# Neoisoptera repetitively colonised Madagascar after the Middle Miocene climatic optimum

**DOI:** 10.1101/2021.12.01.470872

**Authors:** Menglin Wang, Simon Hellemans, Aleš Buček, Taisuke Kanao, Jigyasa Arora, Crystal Clitheroe, Jean-Jacques Rafanomezantsoa, Brian L. Fisher, Rudolf Scheffrahn, David Sillam-Dussès, Yves Roisin, Jan Šobotník, Thomas Bourguignon

**Affiliations:** Okinawa Institute of Science & Technology Graduate University, 1919–1 Tancha, Onna-son, Okinawa, 904–0495, Japan; Faculty of Science, Yamagata University, Yamagata, Japan; Madagascar Biodiversity Center, Parc Botanique et Zoologique de Tsimbazaza, Antananarivo 101, Madagascar; California Academy of Sciences, San Francisco, CA, USA; Fort Lauderdale Research and Education Center, Institute for Food and Agricultural Sciences, 3205 College Avenue, Davie, Florida 33314 USA; Laboratory of Experimental and Comparative Ethology, UR 4443, University Sorbonne Paris Nord, Villetaneuse, France; Faculty of Tropical AgriSciences, Czech University of Life Sciences, Prague, Czech Republic; Evolutionary Biology and Ecology, Université Libre de Bruxelles, Brussels, Belgium

**Keywords:** Australia, endemism, historical biogeography, over-water dispersal, vicariance

## Abstract

Madagascar is home to many endemic plant and animal species owing to its ancient isolation from other landmasses. This unique fauna includes several lineages of termites, a group of insects known for their key role in organic matter decomposition in many terrestrial ecosystems. How and when termites colonised Madagascar remains unknown. In this study, we used 601 mitochondrial genomes, 93 of which were generated from Madagascan samples, to infer the global historical biogeography of Neoisoptera, a lineage containing upwards of 80% of described termite species. Our results indicate that Neoisoptera colonised Madagascar between seven to ten times independently during the Miocene, between 8.4-16.6 Ma (95% HPD: 6.1-19.9 Ma). This timing matches that of the colonization of Australia by Neoisoptera. Furthermore, the taxonomic composition of the Neoisopteran fauna of Madagascar and Australia are strikingly similar, with Madagascar harbouring an additional two lineages absent from Australia. Therefore, akin to Australia, Neoisoptera colonised Madagascar during the global expansion of grasslands, possibly helped by the ecological opportunities arising from the spread of this new biome.

## 1 Introduction

Madagascar is the world’s fourth largest island and is home to a great many endemic plant and animal species [1,2]. One important reason for the peculiarity of its biota is its ancient isolation from other landmasses [2]. Madagascar, together with India, broke away from Africa ∼160 million years ago (Ma) and has retained a distance of ∼400 km from East Africa coast for the last 120 million years [3,4]. India subsequently broke away from Madagascar and started drifting northward, leaving Madagascar separated from other continental landmasses for the last ∼88 Ma [5]. This long isolation is the source of Madagascar’s unique biota.

The fauna of Madagascar has either been interpreted as resulting from vicariance or dispersal origin. Early biogeographers, unaware of the motion of continental landmasses, explained the origin of Madagascar fauna by long-distance over-water dispersals (e.g., Matthew 1915, Simpson 1940) [6,7]. Subsequently, the validation of the continental drift hypothesis [8] in the 1960s initiated a paradigm shift, and vicariance became widely accepted as the dominant mechanism responsible for the Madagascar unique fauna [9,10]. However, the time-calibrated phylogenies produced during the last two decades have revealed that the majority of animal lineages found in Madagascar are younger than the split of Madagascar from continental Africa and India (e.g., Crottini et al. 2012) [11]. This timing implies that Madagascar predominantly acquired its fauna by means of long-distance over-water dispersals after its separation from other landmasses [12]. For instance, the distributions of the extinct elephant bird [13] and the iconic chameleons [14] are explained by such dispersals. Long-distance over-water dispersals also explain the distribution of several Malagasy insect lineages, such as the millipede assassin bugs [15], the beetle tribe Scarabaeini [16], and the hissing cockroaches [17]. Only a few insect lineages, such as the Malagasy alderfly genus *Haplosialis* [18], the cascade beetles [19], and the whirligig beetles [20] are ancient enough to have their modern distribution potentially resulting from vicariance.

Termites are a group of social cockroaches feeding on lignocellulose at various stages of decomposition, from hard wood to the organic matter present in the soil [21]. They include ∼3000 described species mostly distributed across the tropical and subtropical regions [22,23]. The oldest known fossils of termites are ∼130 million years old (Myo) and date back from the early Cretaceous [24,25]. Time-calibrated phylogenies provide slightly older age estimates and suggest that the last common ancestor of modern termites roamed the Earth 140-150 Ma [25–29]. The origin of termites therefore predates the breakup of Gondwana, indicating vicariance may explain the current distribution of early-diverging termite lineages. However, the termite fauna of Madagascar is known to comprise derived genera of Kalotermitidae and Neoisoptera [30–36] and appears to lack early-diverging termite lineages, such as Stolotermitidae and Archotermopsidae, whose distribution may bear the signature of vicariance. Madagascar therefore acquired its modern termite fauna by means of long-distance over-water dispersals, presumably via rafting in floating wood pieces or vegetative rafts that contained parts of termite colonies [37].

The pathways and timing of the spread of termites across continents have been studied in detail in Neoisoptera [38–41]. However, the historical biogeography of Neoisoptera in Madagascar has been largely overlooked. Neoisoptera is composed of four families, Stylotermitidae, Rhinotermitidae, Serritermitidae, and Termitidae, and contains upwards of 80% of described termite species [23,26]. The Neoisoptera are represented in Madagascar by a handful of endemic genera and by a few genera also found in continental Africa and in the Oriental region [30,31,33,35]. The only Malagasy termite lineage whose historical biogeography has been studied in detail is the fungus-growing termite genus *Microtermes*, which colonised Madagascar from continental Africa via a single long-distance over-water dispersal ∼13 Ma [42,43]. This dispersal event was presumably facilitated by the acquisition of a vertical mode of transmission of *Termitomyces* fungal symbionts in *Microtermes* [44]. The timing and geographic origin of other dispersal events, so well as the number of these dispersal events, are presently unknown and require further investigations.

In this study, we reconstructed robust time-calibrated phylogenetic trees of termites using the mitochondrial genomes of 586 Neoisoptera (including 93 Madagascan samples) and 14 outgroups. Our dataset is representative of the worldwide distribution of Neoisoptera and includes species from the Afrotropical, Australian, Madagascan, Nearctic, Neotropical (including Panamanian), Oriental (including Sino-Japanese), Palaearctic, Saharo-Arabian, and Oceanian realms, as defined by Holt et al. (2013) [45]. We used our time-calibrated phylogenetic trees to shed light on the evolution of Neoisoptera, the main termite lineage found in the Madagascan realm. Our specific aims were (i) to provide the first comprehensive phylogenetic tree of Malagasy Neoisoptera; and (ii) to investigate the geographic origin and the timing of dispersal and diversification of the Neoisoptera lineages present in Madagascar.

## 2 Material and Methods

### (a) Biological Samples and Mitochondrial Genome Sequencing

We sequenced the mitochondrial genomes of 92 termite samples from Madagascar. We also sequenced an additional 30 mitochondrial genomes from termite samples collected outside Madagascar, including 13 samples from the Afrotropical realm, two samples from the Saharo-Arabian realm, nine Neotropical samples, one Oceanian sample, and five Nearctic samples (Table S1). These 30 samples mostly belonged to termite lineages present in Madagascar and underrepresented in previous studies, such as *Amitermes, Psammotermes*, and *Prorhinotermes*. Their inclusion improves our reconstructions of ancestral ranges globally and for the Madagascan realm. We combined the 122 mitochondrial genomes sequenced in this study with 478 termite mitochondrial genomes previously published, including the mitochondrial genome of the Madagascan *Prorhinotermes canalifrons* from Reunion Island [27,39–41,46–48]. We also obtained the mitochondrial genome of the cockroach *Cryptocercus relictus* [47], a representative of Cryptocercidae, the sister group of termites. Specimens were tentatively identified based on available taxonomic works and similarity to publicly available COII sequences [23,30,31,36,49,50].

We extracted DNA from two or three individuals preserved in RNA-later® or in 80% ethanol. Samples preserved in RNA-later® were stored at −20°C or −80°C until DNA extraction. Samples preserved in 80% ethanol were stored at room temperature for upwards of 20 years. We used one of the following three sequencing strategies: (i) long-range PCR followed by high-throughput DNA sequencing for samples stored in RNA-later®; (ii) whole-genome shotgun sequencing for samples stored in RNA-later®; and (iii) whole-genome shotgun sequencing for samples stored in 80% ethanol. In all three cases, DNA was extracted with the DNeasy Blood & Tissue extraction kit (Qiagen); and libraries were prepared using the NEBNext® Ultra ™ II FS DNA Library Preparation Kit (New England Biolabs) and the Unique Dual Indexing Kit (New England Biolabs). Libraries were prepared with one-fifteenth of the reagent volumes recommended by the manufacturer.

For the (i) first strategy, DNA was extracted using specimens from which the digestive tract was removed. The whole mitochondrial genomes were amplified in two long-range PCR reactions using the TaKaRa LA Taq polymerase and the primer sets and PCR conditions previously described in Bourguignon et al. (2016)[41]. We mixed both amplicons in equimolar concentration and prepared one library for each sample separately. Libraries were pooled in equimolar concentration and paired-end sequenced using the Illumina Miseq2000 platform. For the (ii) second strategy, whole genomic DNA was extracted from whole body of termite workers including guts. Libraries were pooled in equimolar concentration and paired-end sequenced using the Illumina Hiseq2500 or Hiseq4000 platforms. For the (iii) third strategy, whole genomic DNA was extracted from whole body of termite workers including gut. Libraries were prepared without enzymatic fragmentation step. Libraries were pooled in equimolar concentration and paired-end sequenced using the Illumina HiSeq X or Novaseq platforms.

### (b) Assembly and Alignment

Raw reads were quality-checked with Fastp v0.20.1 [51]. Read adaptors were trimmed. Filtered reads were assembled using metaSPAdes v3.13.0 [52], and retrieved and annotated with MitoFinder v1.4 [53]. IMRA was used as an attempt to elongate mitochondrial genomes that were not assembled in one contig [54]. The control regions were omitted because they present repetitive patterns difficult to assemble with short reads.

All genes were aligned separately. The 22 transfer RNA genes and the two ribosomal RNA genes were aligned as DNA sequences with MAFFT v7.305 [55]. The 13 protein-coding genes were translated into amino acid sequences using EMBOSS v6.6.0 [56]and aligned using MAFFT. Amino acid sequence alignments were back-translated into DNA sequences using Pal2Nal [57]. The 37 gene alignments were concatenated with FASconCAT-G_v1.04.pl [58].

### (c) Phylogenetic Analyses

The concatenated sequence alignment was partitioned into five subsets: one for the combined transfer RNA genes, one for the combined ribosomal RNA genes, and one for each codon position of the protein-coding genes. The phylogenetic analyses were performed with and without third codon positions. Phylogenetic relationships were inferred using maximum likelihood and Bayesian inference methods. We used IQ-TREE v1.6.12 [59] to reconstruct maximum likelihood phylogenetic trees. The best-fit nucleotide substitution model was determined with the Bayesian Information Criterion using ModelFinder [60] implemented in IQ-TREE. Branch supports were estimated using 1000 bootstrap replicates [61]. Bayesian analyses were implemented in MrBayes v3.2.3 using a GTR+G model of nucleotide substitution [62]. The tree and the posterior distribution of parameters were estimated from MCMC samplings. Each analysis was run with four chains, three hot and one cold. Each analysis was run in four replicates to ensure the convergence of the chains. For the analyses with third codon positions included, the chains were run for 40 million generations with a 25% burn-in fraction. For the analyses without third codon positions, the chains were run for 20 million generations with burn-in fraction set to 10%. All the chains were sampled every 5,000 generations. The mixing of the chains and the behaviour of all parameters were examined in Tracer v1.7.1 [63]. For all analyses, the topology was constrained to harbour a sister relationship between the subfamilies Sphaerotermitinae and Macrotermitinae, as supported by transcriptome-based phylogenies [28].

### (d) Divergence time estimation

We analysed the concatenated sequence alignments with and without third codon positions and reconstructed time-calibrated phylogenetic trees using BEAST v2.6.2 [64]. Each analysis was run twice to ensure the convergence of the chain. The rate variation across branches was modelled using an uncorrelated lognormal relaxed clock. We used the Yule model for the tree prior. A GTR+G model of nucleotide substitution was assigned to each partition. For the analyses without third codon positions, we sampled the tree and parameter values of the chain every 50,000 steps over a total of 350 million generations. The first 10% of generations were discarded as burn-in. For the analyses with third codon positions included, the chain was run over 600 million generations and the first 20% sampled trees were discarded. The mixing of the chains and the behaviour of all parameters were examined with Tracer v1.7.1 [63].

The molecular clock was calibrated using 14 fossils as minimum age constraints (Table S2). We used the youngest possible age for each fossil as reported in the Fossilworks database [65] (last accessed on January 2021, 31^st^). We used the criteria described by Parham et al. (2012) to select fossils. For each fossil calibration, we also determined a soft maximum bound using the phylogenetic bracketing approach [66,67]. Each calibration was implemented as exponential priors on node time. We previously justified the use of every fossil calibration used in this study [28]. We used TreeAnnotator to generate a maximum clade credibility consensus tree.

### (e) Biogeographic analyses

We reconstructed the historical biogeography of Neoisoptera using the R package BioGeoBEARS [68]. The Madagascan realm includes Madagascar and neighbouring islands: Comoros, Mascarenes, and Seychelles. We used sampling locations to assign each tip to a biogeographic realm. A total of six phylogenetic reconstructions, estimated with IQ-TREE, MrBayes, and BEAST2 (with and without third codon positions), were subjected to ancestral range reconstructions with BioGeoBEARS. For each phylogenetic tree, we carried out ancestral range reconstructions with the DEC model (Dispersal-Extinction-Cladogenesis), the DIVALIKE model (Dispersal-Vicariance Analysis), and the BAYAREALIKE model. We run each model with and without the parameter “+ j” allowing jump dispersals, which correspond to speciation events following long-distance dispersals [69]. The best-fit model was determined for each phylogenetic reconstruction using AICc (Akaike Information Criterion with sample size corrected).

## 3 Results

### (a) Topology of the phylogenetic trees

Our six phylogenetic trees were largely congruent with respect to interfamily and intergeneric relationships, with the exception of a few nodes with low posterior probabilities and bootstrap supports (Figures 1, S1-6). They showed that the Neoisoptera were represented by species belonging to ten lineages of Rhinotermitidae and Termitidae in the Madagascan realm (Figures 1, S1-S6). The Rhinotermitidae were represented by three species: *Prorhinotermes canalifrons, Coptotermes truncatus*, and *Psammotermes* voeltzkowi. All three species belonged to genera also present in other biogeographic realms. The seven remaining lineages were part of the Termitidae and formed clades endemic to the Madagascan realm, including one clade of Macrotermitinae, two clades of Nasutitermitinae, and four clades of Termitinae. The only Madagascan clade of Macrotermitinae included several species of *Microtermes* that formed the sister group of a clade composed of African *Microtermes* and the Oriental *Ancistrotermes pakistanicus*. One of the two Madagascan clades of Nasutitermitinae contained Malagasitermes milloti, Coarctotermes, and several species assigned to the polyphyletic *Nasutitermes*. The sister group of this clade varied among analyses. The other Madagascan clade of Nasutitermitinae only included *Nasutitermes* sp. 1, retrieved as sister to a group of Oriental species. The four Madagascan clades of Termitinae belonged to *Microcerotermes, Amitermes*, and the *Termes* group, which contained two Madagascan clades. The Madagascan *Microcerotermes* included upwards of nine species with an unresolved sister group. *Amitermes* was represented by two species forming the sister group of a lineage including African, Saharo-Arabian, Oriental, and Australian *Amitermes*. One of the two Madagascan clades of the *Termes* group included *Quasitermes, Capritermes*, and a species resembling *Quasitermes*. This first clade was sister to a clade containing the Malagasy and Oriental species of *Termes* as well as the Australian members of the *Termes* group.

**Figure 1.**
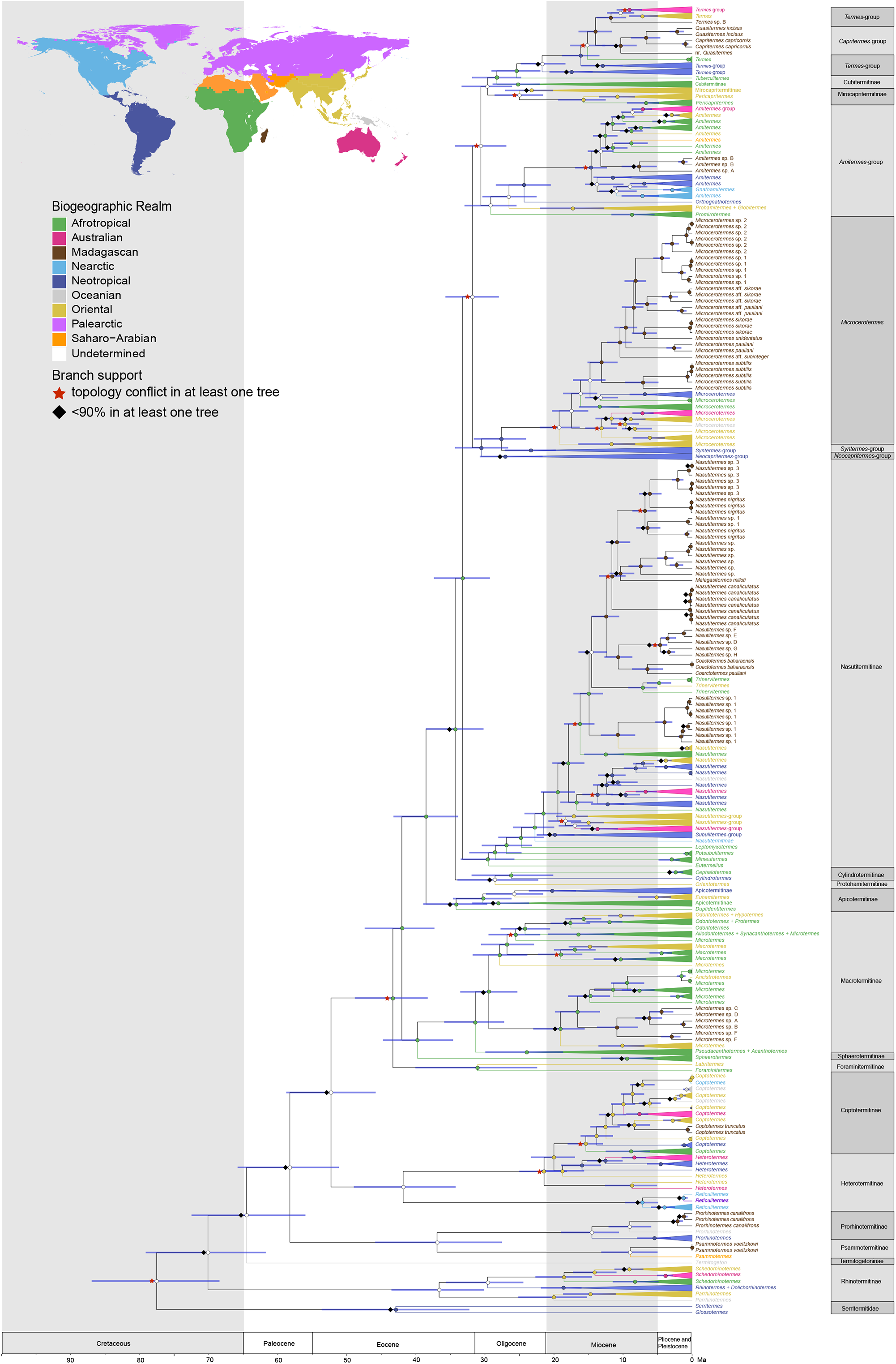
Time-calibrated Bayesian phylogeny inferred from 601 mitochondrial genomes, with the third codon position excluded. Node labels provide a summary of node supports across the six phylogenetic analyses: red stars indicate topology conflicts for at least one analysis; and black diamond indicate support value <90% for at least one analysis. Nodes without labels have support values >90% for all analyses. Node bars indicate the 95% HPD intervals estimated with BEAST 2. Tips and node circles are color-coded to indicate biogeographic realms. The colours of node circles indicate ancestral ranges reconstructed with probabilities higher than 65% for the six phylogenetic trees inferred with IQ-TREE, MrBayes, and BEAST 2, with and without third codon positions. White circles indicate undetermined ancestral distribution. Clades containing species collected in the same biogeographic realm are collapsed, except for species collected in the Madagascan realm. The map shows the biogeographic realms recognised in this study (modified from Holt et al. 2013).

### (b) Divergence times

The time-calibrated phylogenetic trees reconstructed with and without third codon positions of protein-coding genes diverged in their age estimates by up to 5 million years (Figures 1, S1-2). The divergences were smaller than 2.2 million years for the nodes representing the splits between Madagascan clades and their sister groups (Figures 1, S1-2). Given the similar divergence age estimates obtained with both analyses, we will only discuss the results of the analysis with third codon positions excluded for the sake of simplicity.

All Madagascan clades of Neoisoptera diverged from their sister groups during the Miocene (Figure 2). Within the Rhinotermitidae, the Madagascan *Prorhinotermes, Psammotermes*, and *Coptotermes* diverged from their sister groups 9.0 Ma (95% height posterior density (HPD): 6.0-12.1 Ma), 9.0 Ma (95% HPD: 5.0-13.2 Ma), and 8.4 Ma (95% HPD: 6.1-10.8 Ma), respectively. Within the Termitidae, the Madagascan macrotermitine *Microtermes* diverged from their sister group 16.6 Ma (95% HPD: 13.4-19.9 Ma). The Madagascan nasutitermitine clade containing Malagasitermes milloti and Coarctotermes diverged from its sister 14.4 Ma (95% HPD: 12.3-16.5 Ma). The other Madagascan nasutitermitine clade, composed of *Nasutitermes* sp. 1, diverged from its Oriental sister group 10.7 Ma (95% HPD: 8.2-13.2 Ma). Within the termitines, we dated the divergence between the Madagascan *Amitermes* and other *Amitermes* species at 13.1 Ma (95% HPD: 11.3-15.1 Ma). The most recent common ancestor of all Madagascan *Microcerotermes* and their sister group was estimated at 14.8 Ma (95% HPD: 12.5-17.2 Ma). The Madagascan *Quasitermes* + *Capritermes* clade diverged from its sister group 14.0 Ma (95% HPD: 11.6-16.5 Ma), while the Madagascan *Termes* sp. B diverged from its sister group 11.7 Ma (95% HPD: 9.5-13.9 Ma).

**Figure 2.**
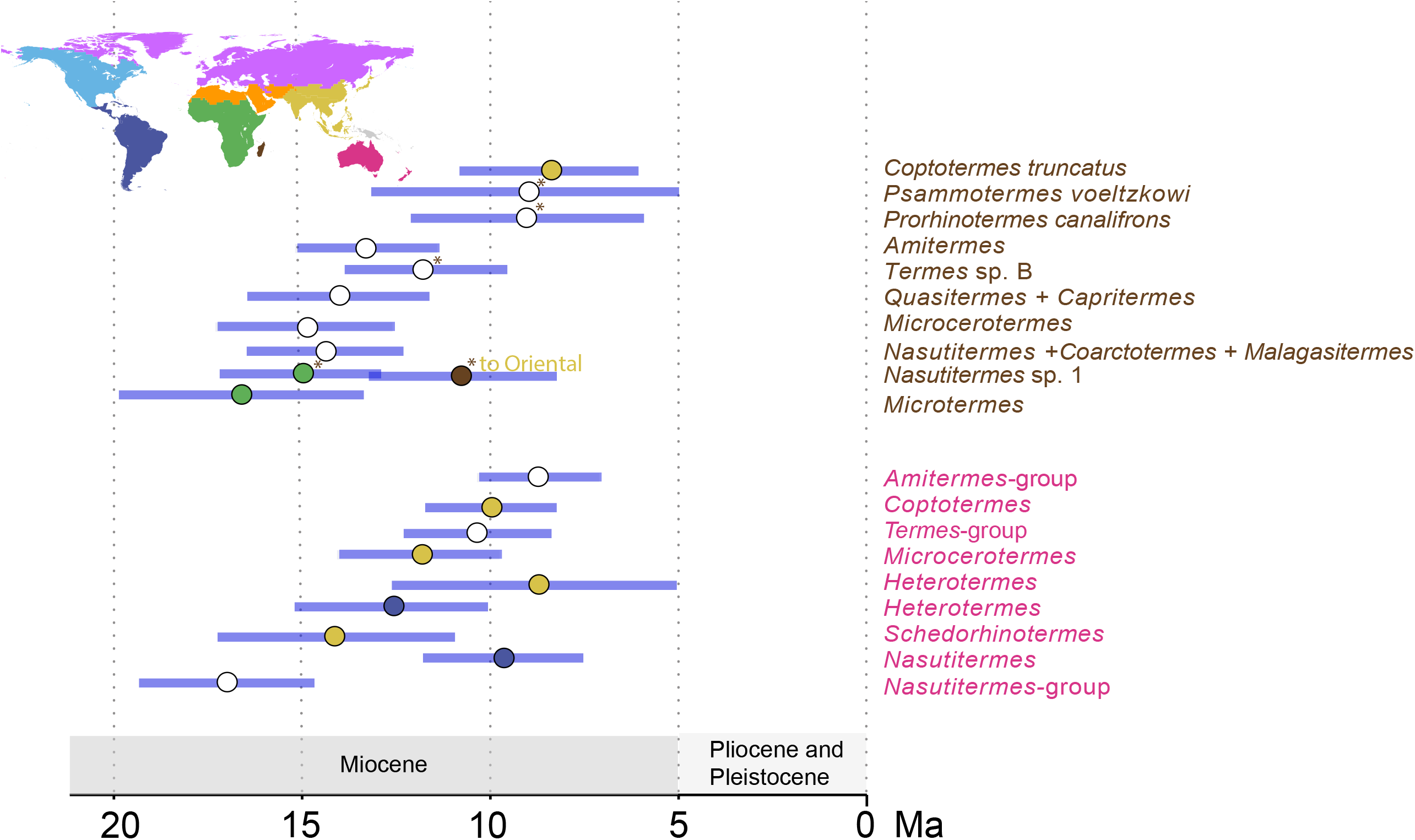
Summary of Madagascar and Australia colonization events. The scenario with 10 dispersal events is displayed. Node bars indicate the 95% HPD intervals estimated with BEAST 2. The colours of node circles indicate ancestral ranges reconstructed with probabilities higher than 65% for the six phylogenetic trees inferred with IQ-TREE, MrBayes, and BEAST 2, with and without third codon positions. White circles indicate undetermined ancestral distribution. The map shows the biogeographic realms recognised in this study (modified from Holt et al. 2013). Asterisks indicate ambiguous dispersal events.

### (c) Biogeographic reconstruction

We reconstructed the ancestral range distribution of Neoisoptera on our six phylogenetic trees using six different models. The DEC + *j* model was the best-fit model for all trees, except for the BI tree without third positions for which the best-fit model was the DIVALIKE + *j* model (for details, see Table S3). The models with the parameter + *j* fit the data better than the models without this parameter, indicating that jump dispersals played a major role in the biogeographic history of Neoisoptera.

Our analyses indicated that the Madagascan realm was colonised by seven to ten long-distance over-water dispersals (Figure 2). Four Neoisopteran lineages unambiguously colonised the Madagascan realm once: *Coptotermes truncatus* colonised the Madagascan realm from the Oriental realm; *Microtermes* from the Afrotropical realm; and *Microcerotermes* and *Amitermes* from an unidentified realm. The colonization of the Madagascan realm by *Prorhinotermes* + *Psammotermes*, the Nasutitermitinae, and the *Termes* group involved one or two long-distance over-water dispersals. Following the most likely scenario, *Prorhinotermes* and *Psammotermes* independently colonised the Madagascan realm through long-distance over-water dispersals from undetermined biogeographic realms. The alternative scenario of an early arrival of the common ancestor of *Prorhinotermes* and *Psammotermes*, followed by subsequent long-distance over-water dispersals to other biogeographic realms, was less likely but could not be excluded. Similarly, the two Madagascan clades of Nasutitermitinae probably originated from two independent dispersals from Africa to the Madagascan realm. A less likely alternative featured one long-distance over-water dispersal from Africa to the Madagascan realm followed by two dispersal events from Madagascar to the Afrotropical and Oriental realms. Lastly, the Madagascan realm was either colonised once by the *Termes* group followed by one or several dispersal events out of the Madagascan realm, or it was independently colonised twice, once by each Malagasy lineage of the *Termes* group.

## 4 Discussion

### (a) Long-distance over-water dispersals of Neoisoptera to and from the Madagascan realm: taxonomic identity, timing, and origin of the dispersers

We reconstructed the most comprehensive phylogenetic tree of Neoisoptera to date. The relationships among the main lineages of Neoisoptera were largely congruent with earlier molecular studies based on mitochondrial genome and transcriptome data [27,28,40]. Our time estimates were generally younger than those found by these studies, but remained congruent, with overlapping HPD intervals. These differences may pertain to the use of different fossil calibrations and/or to the changes of fossil age estimations at the time of publication of these studies. For example, we previously used the ∼110 Myo *Cratokalotermes santanensis* [70] to calibrate Kalotermitidae + Neoisoptera [27,40] while we now use the ∼95 Myo *Archeorhinotermes rossi* [71] to calibrate the same node.

Before this study, three genera of Rhinotermitidae —*Coptotermes, Prorhinotermes*, and *Psammotermes*— and four groups of Termitidae —*Microtermes, Microcerotermes*, the Nasutitermitinae, and the *Termes* group— were known to be represented by species native to the Madagascan realm[23,30,31,33,35,72]. We sequenced upward of 40 Malagasy species, while Eggleton and Davies (2003) listed 33 species of Neoisoptera described from Madagascar, implying the existence of several new species among our samples [72]. The most notable species were two new species of *Amitermes*, a genus previously unknown from the Madagascan realm. We also found that the Malagasy species of Nasutitermitinae and of the *Termes* group do not form monophyletic groups. Therefore, the Madagascan termite fauna is more phylogenetically diverse than previously envisioned.

Our ancestral state reconstructions indicated that Neoisoptera colonised the Madagascan realm seven to ten times independently and possibly dispersed out of the Madagascan realm up to four times. The dispersal events to and from the Madagascan realm took place 8.4-16.6 Ma (95% HPD: 6.1-19.9 Ma), between the mid-Miocene climatic optimum [73] and the end of the Miocene. Therefore, our results indicate that the Madagascan realm acquired its fauna of Neoisoptera through long-distance over-water dispersal events.

Our ancestral range reconstructions also revealed one long-distance over-water dispersal event within the Madagascan realm, that of *Prorhinotermes canalifrons* between Madagascar and the Reunion Island 2.1 Ma (95% HPD: 1.3-3.0 Ma). This species is also known from Mauritius, Comoros, and Seychelles [23], potentially indicating additional over-water dispersals among islands of the Madagascan realm for this genus with high dispersal abilities and tolerance to salinity [74,75]. Two other species, *Coptotermes truncatus* and *Microcerotermes subtilis*, as well as the nasutitermitine genus *Kaudernitermes*, are also known from Madagascar and several neighbouring islands [23], indicating further dispersals between islands. Whether these dispersals were mediated by human activities or were long-distance over-water dispersals, as was the case for P. *canalifrons* in Madagascar and the Reunion islands, is unclear. Additional sequence data from the termite fauna of the Reunion Island, Mauritius, Comoros, and Seychelles are needed to identify the processes of colonization of these islands.

We were able to identify the source of three dispersal events to the Madagascan realm: *Coptotermes truncatus* has Oriental origin, *Microtermes* has African origin [42], and the nasutitermitines arrived from Africa at least once. The origin of other Madagascan lineages remains unresolved. Therefore, our results do not provide a global picture of the origin of the Madagascan Neoisoptera, although we show that some lineages have African and Oriental origins, as is the case for many other taxa [12,76]. Our ancestral range reconstructions also point to the possibility that Neoisoptera dispersed out of the Madagascan realm on multiple occasions, although these events remain speculative. Additional sequences from other biogeographical realms are required to identify the origin of Madagascan Neoisoptera lineages with yet unresolved origin.

### (b) The colonization of the Madagascan and Australian realms by Neoisoptera coincides with the global expansion of grasslands

The colonization of the Madagascan realm by Neoisoptera coincides with the colonization of the Australian realm [39–41,77] (Figure 2). The colonization of both realms was initiated around the Miocene climatic optimum, 15-17 Ma, and continued over the next five to ten million years while the world climate gradually cooled down [73] and grasslands expanded worldwide [78]. It is therefore tempting to attribute this coincidental timing to shared historical climatic and ecological changes in Australia and Madagascar.

The climate of Australia became drier during the Middle Miocene ∼14 Ma, and new biomes composed of flora and fauna adapted to arid conditions expended [79,80]. The expansion of the arid-adapted biomes in Australia was accompanied by the opening of new ecological opportunities for local Australian taxa and for colonisers arriving from other continents [79], which included a dozen of lineages of Neoisoptera [39–41,77] (Figure 2). Unlike in Australia, the origin of grasslands in Madagascar is still debated. Human activities have undoubtedly contributed to the expansion of modern Madagascar’s grasslands, and some authors have argued that, prior to human arrival, the areas presently covered by grasslands were forested and only contained patches of grasslands [81,82]. The alternative view is that Madagascar’s grassland first appeared during the Miocene and gradually expanded, an expansion that was accelerated by human arrival [83–85]. Whichever scenario turns out to be correct, the arrival of Neoisoptera in the Madagascan realm was concurrent to the diversification of grasses in Madagascar, whose number of species exponentially increased since around 20 Ma [86]. The divergence between the two grass-feeding species *Coarctotermes pauliani* and *Coarctotermes baharaensis* 6.5 Ma (95% HPD: 4.2-8.8 Ma) indicates an early adaptation of some termite species to grassland in Madagascar. However, the bulk of the termite diversity in Madagascar is associated with forested areas [72]. The colonization of Madagascar and Australia by Neoisoptera therefore coincides with the global spread of grasses.

In addition to the timing of colonization, another parallel that can be made between the Neoisopteran fauna of the Madagascan and Australian realms is the similarity of their taxonomic composition. The Madagascan realm was colonised by two genera not found in Australia, the rhinotermitid *Psammotermes* and the termitid *Microtermes* [30], while the Australian realm was colonised by three genera absent from Madagascar, the rhinotermitids *Schedorhinotermes, Parrhinotermes*, and *Heterotermes* [87]. Note that *Heterotermes* philippinensis was introduced in Madagascar and in Mauritius [88,89]. In contrast, both realms were colonised by the rhinotermitid genera *Coptotermes* and *Prorhinotermes* and by the termitid genera *Microcerotermes, Amitermes, Termes*, and *Nasutitermes* [30,87]. Of note, the latter three genera are paraphyletic and include a number of genera endemic to the Madagascan and Australian realms nested within them. Therefore, these two realms host taxonomically similar communities of Neoisoptera, acquired within the same geological time interval. These observations suggest the existence of ecological preadaptations in the Neoisopteran lineages that colonised Madagascar and Australia, two distant landmasses presently dominated by grasslands and savannah biomes.

We previously reconstructed the global spread of Neoisoptera without samples from Madagascar [38–41]. The sequencing of 92 termite samples from Madagascar provides an opportunity to refine the picture of the global spread of Neoisoptera. The higher termites, which make up over 80% of species of Neoisoptera [23], originated from Africa and dispersed worldwide in two phases [40]. During the first phase, which spanned the Oligocene and the early Miocene, ∼34-20 Ma, higher termites colonised the Neotropical and Oriental realms via a dozen of over-water dispersal events [40]. Our results show that the second phase, which took place during the Miocene, ∼20-8 Ma, was characterised by the colonization of Australia and Madagascar by higher termites and coincides with the global expansion of grasslands.

## Acknowledgments

We thank the DNA Sequencing Section and the Scientific Computation and Data Analysis Section (SCDA) of the Okinawa Institute of Science and Technology Graduate University, Okinawa, Japan, for assistance with sequencing and for providing access to the OIST computing cluster, respectively. This work was supported by the subsidiary funding to OIST, by the Czech Science Foundation (project No. 15-07015Y), by the Internal Grant Agency of the Faculty of Tropical AgriSciences, CULS (20213112), by the Japan Society for the Promotion of Science through a DC2 graduate student fellowship to JA and a postdoctoral fellowship to SH (19F19819).

