## Supplementary legends for "Neoisoptera repetitively colonised Madagascar after the Middle Miocene climatic optimum"

**Supplementary Materials:**

Figure S1: Time-calibrated phylogenetic tree of Neoisoptera reconstructed with BEAST2 using mitochondrial genomes without third codon positions. The ancestral range reconstruction was performed with the best-fit model (see Table 3): the DIVALIKE+J model of BioGeoBEARS. Node bars indicate the 95% HPD intervals. Posterior probabilities are indicated for nodes with support < 0.9. Pie charts on the nodes show the reconstructed ancestral ranges. Colors represent the biogeographical realms recognized in this study (see Holt et al. 2013).

Figure S2: Time-calibrated phylogenetic tree of Neoisoptera reconstructed with BEAST2 using mitochondrial genomes with third codon positions included. The ancestral range reconstruction was performed with the best-fit model (see Table 3): the DEC+J model of BioGeoBEARS. Node bars indicate the 95% HPD intervals. Posterior probabilities are indicated for nodes with support < 0.9. Pie charts on the nodes show the reconstructed ancestral ranges. Colors represent the biogeographical realms recognized in this study (see Holt et al. 2013).

Figure S3: Maximum likelihood phylogenetic tree of Neoisoptera reconstructed with IQ-TREE using mitochondrial genomes without third codon positions. The ancestral range reconstruction was performed with the best-fit model (see Table 3): the DEC+J model of BioGeoBEARS. Node bars indicate the 95% HPD intervals. Posterior probabilities are indicated for nodes with support < 0.9. Pie charts on the nodes show the reconstructed ancestral ranges. Colors represent the biogeographical realms recognized in this study (see Holt et al. 2013).

Figure S4: Maximum likelihood phylogenetic tree of Neoisoptera reconstructed with IQ-TREE using mitochondrial genomes with third codon positions included. The ancestral range reconstruction was performed with the best-fit model (see Table 3): the DEC+J model of BioGeoBEARS. Node bars indicate the 95% HPD intervals. Posterior probabilities are indicated for nodes with support < 0.9. Pie charts on the nodes show the reconstructed ancestral ranges. Colors represent the biogeographical realms recognized in this study (see Holt et al. 2013).

Figure S5: Bayesian phylogenetic tree of Neoisoptera reconstructed with MrBayes using mitochondrial genomes without third codon positions. The ancestral range reconstruction was performed with the best-fit model (see Table 3): the DIVALIKE+J model of BioGeoBEARS. Node bars indicate the 95% HPD intervals. Posterior probabilities are indicated for nodes with support < 0.9. Stars indicate unresolved polytomies in the original MrBayes tree. Pie charts on the nodes show the reconstructed ancestral ranges. Colors represent the biogeographical realms recognized in this study (see Holt et al. 2013).

Figure S6: Bayesian phylogenetic tree of Neoisoptera reconstructed with MrBayes using mitochondrial genomes with third codon positions included. The ancestral range reconstruction was performed with the best-fit model (see Table 3): the DEC+J model of BioGeoBEARS. Node bars indicate the 95% HPD intervals. Posterior probabilities are indicated for nodes with support < 0.9. Stars indicate unresolved polytomies in the original MrBayes tree. Pie charts on the nodes show the reconstructed ancestral ranges. Colors represent the biogeographical realms recognized in this study (see Holt et al. 2013).

Table S1: Samples used in this study, with corresponding collection details and accession numbers.

Table S2: Fossils used for time calibrations in this study.

Table S3: Statistical comparisons of DEC, DEC+J, DIVALIKE, DIVALIKE+J, BAYAREALIKE and BAYAREALIKE+J for the six phylogenetic trees reconstructed in this study. Likelihood Ratio Tests (LRT) were performed to determine the effect of jumping speciation events for each pair of models. Models were statistically compared using corrected Aikake Information Criterion (AICc) weights. Abbreviations: *d*, dispersal rate between biogeographic realms; *e*, local extinction rate.
